# Gene model for the ortholog of *Ilp4* in *Drosophila eugracilis*

**DOI:** 10.1101/2025.09.05.674564

**Authors:** Rachael Cowan, Amun Uppal, Clairine I. S. Larsen, Christopher E. Ellison, Nicole S. Torosin, Jeffrey S. Thompson, Chinmay P. Rele, Lori Boies

**Affiliations:** The University of Alabama, Tuscaloosa, AL, USA; Rutgers University, New Brunswick, NJ USA; Denison University, Granville, OH, USA; St. Mary’s University, San Antonio, TX, USA

## Abstract

Gene Model for *Insulin-like peptide 4* (*Ilp4)* in the *D. eugracilis* (DeugGB2) assembly (GCA_000236325.2). The characterization of this ortholog was carried out as part of a larger, ongoing dataset designed to explore the evolution of the insulin/insulin-like growth factor signaling (IIS) pathway across the genus *Drosophila*, utilizing the Genomics Education Partnership gene annotation protocol within Course-based Undergraduate Research Experiences.

## Introduction

This article reports a predicted gene model generated by undergraduate work using a structured gene model annotation protocol defined by the Genomics Education Partnership (GEP; thegep.org) for Course-based Undergraduate Research Experience (CURE). The following information in quotes may be repeated in other articles submitted by participants using the same GEP CURE protocol for annotating Drosophila species orthologs of Drosophila melanogaster genes in the insulin signaling pathway.

There are eight insulin-like peptides in *Drosophila* which act on the insulin receptor to trigger the Insulin-like Receptor (IR) signaling pathway (Guirao-Rico *et al*., 2010; Brogiolo *et al*., 2001). This family of peptides affects growth and metabolism in *Drosophila* and has been found to impact female sleep and mating behavior (Wu and Brown, 2005; Wigby *et al*., 2011; Newell *et al*., 2020). *Ilp4* is highly expressed in embryonic mesoderm and midgut and larval midgut (Brogiolo *et al*., 2001). Grönke *et al*. found that out of the insulin-like peptides 1-7, *Ilp4* is the most conserved between *Drosophila* species after *Ilp7* (2010). Despite this conservation, little is known about the specific function of Ilp4.

We propose a gene model for the *D. eugracilis* ortholog of the *D. melanogaster Insulin-like peptide 4 (Ilp4)* gene. The genomic region of the ortholog corresponds to the uncharacterized protein LOC108114483 (RefSeq accession XP_017080990.1) in the Deug_2.0 Genome Assembly of *D. eugracilis* (GenBank Accession: GCA_000236325.2 - Chen et al., 2014). This model is based on RNA-Seq data from *D. eugracilis* (PRJNA63469) and *Ilp4* in *D. melanogaster* using FlyBase release FB2022_04 (GCA_000001215.4; Larkin et al., 2021). *D. eugracilis* is part of the *melanogaster* species group within the subgenus *Sophophora* of the genus *Drosophila* (Pélandakis et al., 1993). It was first described as *Tanygastrella gracilis* by Duda (1924) and revised *to Drosophila eugracilis* by Bock and Wheeler (1972). *D. eugracilis* is found in humid tropical and subtropical forests across southeast Asia (Morgan et al., 2022).

“In this GEP CURE protocol students use web-based tools to manually annotate genes in non-model Drosophila species based on orthology to genes in the well-annotated model organism, the fruit fly Drosophila melanogaster. This allows undergraduates to participate in course-based research by generating manual annotations of genes in non-model species (Rele et al., 2023). Computational-based gene predictions in any organism are often improved by careful manual annotation and curation, allowing for more accurate analyses of gene and genome evolution (Mudge and Harrow, 2016; Tello-Ruiz et al., 2019). These models of orthologous genes across species, such as the one presented here, then provide a reliable basis for further evolutionary genomic analyses when made available to the scientific community.” (Myers et al., 2024).

## Results

### Synteny

The target gene, *Ilp4*, occurs on chromosome 3L in *D. melanogaster* is nested within *CG32052* alongside Insulin-like peptide 3 (*Ilp3*) and Insulin-like peptide 2 (*Ilp2*). *Ilp4* is flanked upstream by Cyclin-dependent kinase 8 (*Cdk8*), Inhibitor-2 (*I-2*), and Insulin-like peptide 5 (Ilp5) which is nested by *CG43897. Ilp4* is flanked downstream by Insulin-like peptide 1 (*Ilp1*) and Z band alternatively spliced PDZ-motif protein 67 (*Zasp67*). The *tblastn* search of *D. melanogaster* Ilp4-PA (query) against the *D. eugracilis* (GenBank Accession: GCA_000236325.2) Genome Assembly (database) placed the putative ortholog of *Ilp4* within scaffold scf7180000409711 (KB465257.1) at locus LOC108114483 (XP_017080990.1)— with an E-value of 8e-23 and a percent identity of 43.94%. Furthermore, the putative ortholog nested within LOC108114479 (XP_017080986.1) alongside LOC108114482 (XP_017080989.1), LOC108113894 (XP_017080088.1), and LOC108114481 (XP_017080988.1), which correspond to *CG32052, Ilp3, CG33483*, and *Ilp2* in *D. melanogaster* (E-value: 0.0, 2e-48, 1e-68, and 3e-53; identity: 91.26%, 68.63%, 57.40%, and 59.71%, respectively, as determined by *blastp*; Figure 1A, Altschul et al., 1990). The putative ortholog of *Ilp4* is flanked upstream by LOC108114394 (XP_017080854.1), LOC108114226 (XP_017080582.1), and LOC108114228 (XP_017080583.1) which is nested by LOC108114224 (XP_017080567.1), which correspond to *Cdk8, I-2, Ilp5*, and *CG43897* in *D. melanogaster* (E-value: 0.0, 1e-100, 2e-38, and 0.0; identity: 97.14%, 85.37%, 56.25%, and 80.83%, respectively, as determined by *blastp*). The putative ortholog of *Ilp4* is flanked downstream by LOC108114480 (XP_017080987.1) and LOC108114478 (XP_017080982.1), which correspond to *Ilp1* and *Zasp67* in *D. melanogaster* (E-value: 3e-59 and 0.0; identity: 63.64% and 82.66%, respectively, as determined by *blastp*). The putative ortholog assignment for *Ilp4* in *D. eugracilis* is supported by the following evidence: The genes surrounding the *Ilp4* ortholog are orthologous to the genes at the same locus in *D. melanogaster* and synteny is completely conserved, supported by e-values and percent identities, so we conclude that LOC108114483 is the correct ortholog of *Ilp4* in *D. eugracilis* (Figure 1A).

**Figure 1:**
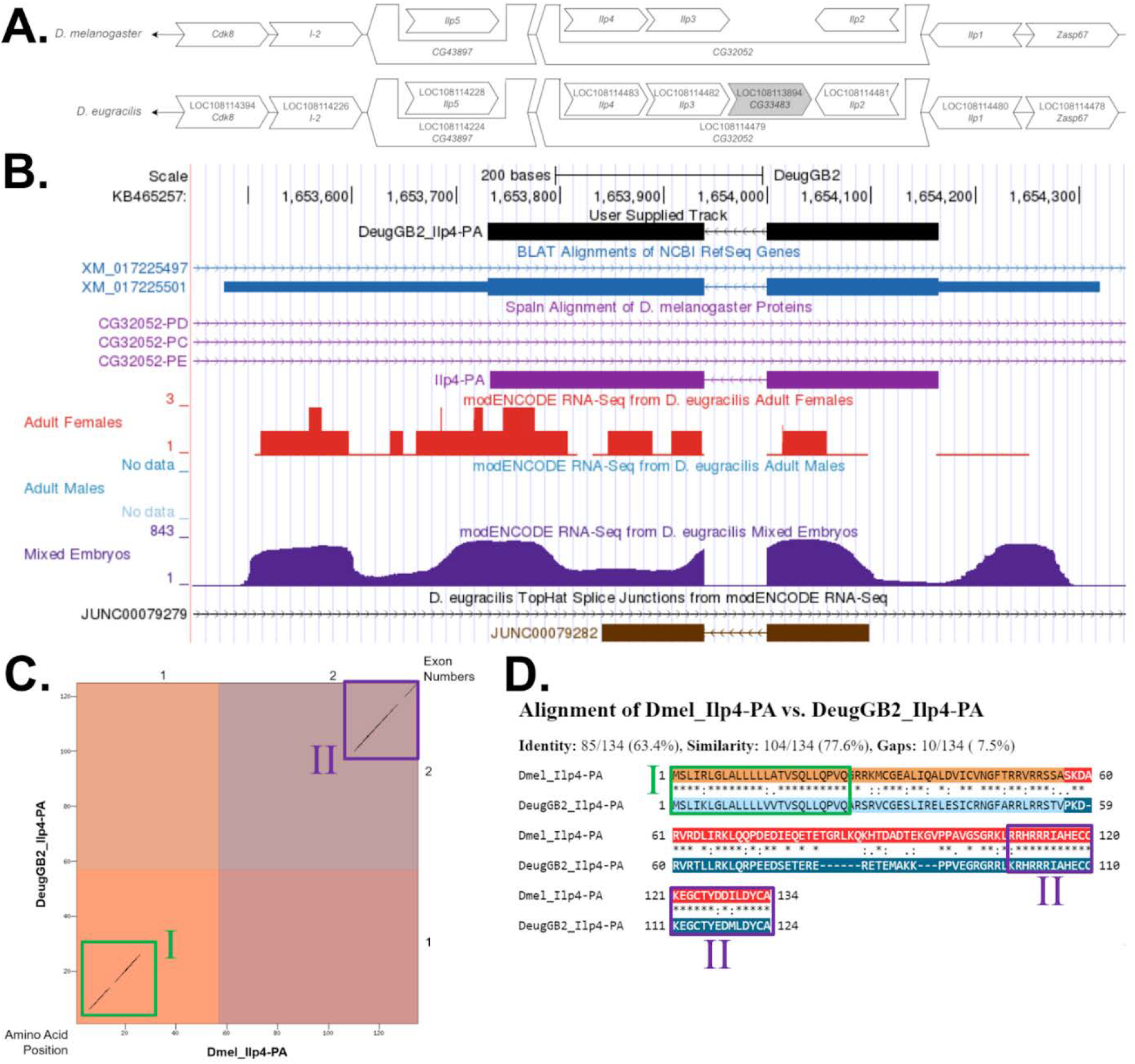
Ilp4 gene model comparison between Drosophila eugracilis and Drosophila melanogaster. **(A) Synteny comparison of the genomic neighborhoods for *Ilp4* in *Drosophila melanogaster* and *D. eugracilis***. Thin underlying arrows indicate the DNA strand within which the target gene–*Ilp4*–is located in *D. melanogaster* (top) and *D. eugracilis* (bottom). The thin arrows pointing to the left indicate that *Ilp4* is on the negative (-) strand in *D. eugracilis* and *D. melanogaster*. The wide gene arrows pointing in the same direction as *Ilp4* are on the same strand relative to the thin underlying arrows, while wide gene arrows pointing in the opposite direction of *Ilp4* are on the opposite strand relative to the thin underlying arrows. White gene arrows in *D. eugracilis* indicate orthology to the corresponding gene in *D. melanogaster*, while grey gene arrows indicate a gene insertion in *D. eugracilis*. Gene symbols given in the *D. eugracilis* gene arrows indicate the orthologous gene in *D. melanogaster*, while the locus identifiers are specific to *D. eugracilis*. **(B) Gene Model in GEP UCSC Track Data Hub (Raney et al**., **2014)**. The coding-regions of *Ilp4* in *D. eugracilis* are displayed in the User Supplied Track (black); coding exons are depicted by thick rectangles and introns by thin lines with arrows indicating the direction of transcription. Subsequent evidence tracks include BLAT Alignments of NCBI RefSeq Genes (dark blue, alignment of Ref-Seq genes for *D. eugracilis*), Spaln of D. melanogaster Proteins (light purple, alignment of Ref-Seq proteins from *D. melanogaster*), RNA-Seq from Adult Females and Mixed Embryos (red and dark purple, respectively; alignment of Illumina RNA-Seq reads from *D. eugracilis*), and Splice Junctions Predicted by regtools using *D. eugracilis* RNA-Seq (PRJNA63469). The splice junction which aligns with *Ilp4* (JUNC00079282) has a read-depth of 657. **(C) Dot Plot of Ilp4-PA in *D. melanogaster* (*x*-axis) vs. the orthologous peptide in *D. eugracilis* (*y*-axis)**. Amino acid number is indicated along the left and bottom; coding-exon number is indicated along the top and right, and exons are also highlighted with alternating colors. The black diagonal lines indicate sequence similarity and are highlighted by the green box denoted I and the purple box denoted II. **(D) Protein alignment between *D. melanogaster* Ilp4-PA and its putative ortholog in *D. eugracilis***. The alternating colored rectangles represent adjacent exons. The symbols in the match line denote the level of similarity between the aligned residues. An asterisk (^*^) indicates that the aligned residues are identical. A colon (:) indicates the aligned residues have highly similar chemical properties—roughly equivalent to scoring > 0.5 in the Gonnet PAM 250 matrix (Gonnet et al., 1992). A period (.) indicates that the aligned residues have weakly similar chemically properties—roughly equivalent to scoring > 0 and ≤ 0.5 in the Gonnet PAM 250 matrix. A space indicates a gap or mismatch when the aligned residues have a complete lack of similarity—roughly equivalent to scoring ≤ 0 in the Gonnet PAM 250 matrix. The green box denoted I and the purple boxes denoted II correspond to the boxes in figure 1C, respectively.

### Protein Model

*Ilp4* in *D. eugracilis* has one protein-coding isoform (Ilp4-PA; Figure 1B). Isoform (Ilp4-PA) contains two protein-coding exons. Relative to the ortholog in *D. melanogaster*, the coding-exon number and isoform count are conserved. The sequence of Ilp4-PA in *D. eugracilis* has 60.00% identity (E-value: 1e-38) with the protein-coding isoform Ilp4-PA in *D. melanogaster*, as determined by *blastp* (Figure 1C). Unusual characteristics of this model include the low sequence similarity across the end of protein-coding exon one and beginning of protein-coding exon two, spanning the region in between green box I and purple box(es) II (Figures 1C, 1D).

### RNA-Seq Data

The RNA-Seq data supported the final reconciled model. Peaks in the data correspond to exons while low coverage areas indicate introns. There was no RNA-Seq data for adult males, and the embryonic data reports higher reads than the adult female track which has very low coverage. *Brogiolo et al*. found that *Ilp4* is highly expressed in embryonic mesoderm and midgut, and the RNA-Seq data supports the higher expression of *Ilp4* in *Drosophila* embryos. Any RNA-Seq reads that extend past the 5’ and 3’ ends of the model can be attributed to untranslated regions and to the nesting gene.

## Methods

“Detailed methods including algorithms, database versions, and citations for the complete annotation process can be found in Rele et al. (2023). Briefly, students use the GEP instance of the UCSC Genome Browser v.435 (https://gander.wustl.edu; Kent WJ et al., 2002; Navarro Gonzalez et al., 2021) to examine the genomic neighborhood of their reference IIS gene in the *D. melanogaster* genome assembly (Aug. 2014; BDGP Release 6 + ISO1 MT/dm6). Students then retrieve the protein sequence for the *D. melanogaster* reference gene for a given isoform and run it using *tblastn* against their target *Drosophila* species genome assembly on the NCBI BLAST server (https://blast.ncbi.nlm.nih.gov/Blast.cgi; Altschul et al., 1990) to identify potential orthologs. To validate the potential ortholog, students compare the local genomic neighborhood of their potential ortholog with the genomic neighborhood of their reference gene in *D. melanogaster*. This local synteny analysis includes at minimum the two upstream and downstream genes relative to their putative ortholog. They also explore other sets of genomic evidence using multiple alignment tracks in the Genome Browser, including BLAT alignments of RefSeq Genes, Spaln alignment of *D. melanogaster* proteins, multiple gene prediction tracks (e.g., GeMoMa, Geneid, Augustus), and modENCODE RNA-Seq from the target species. Detailed explanation of how these lines of genomic evidenced are leveraged by students in gene model development are described in Rele et al. (2023). Genomic structure information (e.g., CDSs, intron-exon number and boundaries, number of isoforms) for the *D. melanogaster* reference gene is retrieved through the Gene Record Finder (https://gander.wustl.edu/~wilson/dmelgenerecord/index.html; Rele et al., 2023). Approximate splice sites within the target gene are determined using *tblastn* using the CDSs from the *D. melanogaste*r reference gene. Coordinates of CDSs are then refined by examining aligned modENCODE RNA-Seq data, and by applying paradigms of molecular biology such as identifying canonical splice site sequences and ensuring the maintenance of an open reading frame across hypothesized splice sites. Students then confirm the biological validity of their target gene model using the Gene Model Checker (https://gander.wustl.edu/~wilson/dmelgenerecord/index.html; Rele et al., 2023), which compares the structure and translated sequence from their hypothesized target gene model against the *D. melanogaster* reference gene model. At least two independent models for a gene are generated by students under mentorship of their faculty course instructors. Those models are then reconciled by a third independent researcher mentored by the project leaders to produce the final model. Note: comparison of 5’ and 3’ UTR sequence information is not included in this GEP CURE protocol.” (Gruys et al, 2025)

## Supporting information

FASTA, GFF, PEP Files

## Supplemental Material

1. Zip file containing FASTA, PEP, GFF files for the gene model
2. Figure 1 in high resolution

### Metadata

Bioinformatics, Genomics, *Drosophila*, Genotype Data, New Finding

## Acknowledgements

We thank Wilson Leung (Washington University, St. Louis) for developing and maintaining the technological infrastructure that supported the creation of this gene model, as well as Chinmay Rele and Laura K. Reed (University of Alabama) for their guidance and encouragement throughout the project. We are also grateful to FlyBase for providing the authoritative database for *Drosophila melanogaster* gene models. FlyBase is supported by grants NHGRI U41HG000739 and U24HG010859, UK Medical Research Council MR/W024233/1, NSF 2035515 and 2039324, BBSRC BB/T014008/1, and Wellcome Trust PLM13398.

## Funding

This material is based upon work supported by the National Science Foundation (1915544) and the National Institute of General Medical Sciences of the National Institutes of Health (R25GM130517) to the Genomics Education Partnership (GEP; https://thegep.org/; PI-Laura K. Reed). Any opinions, findings, and conclusions or recommendations expressed in this material are solely those of the author(s) and do not necessarily reflect the official views of the National Science Foundation nor the National Institutes of Health.

## Notes

### Competing Interest Statement

The authors have declared no competing interest.

